# Computational Insights into Membrane Disruption by Cell-Penetrating Peptides

**DOI:** 10.1101/2024.10.09.617374

**Authors:** Eric Catalina-Hernandez, Marcel Aguilella-Arzo, Alex Peralvarez-Marin, Mario Lopez-Martin

**Affiliations:** Unit of Biophysics, Department of Biochemistry and Molecular Biology, Facultat de Medicina, Av. Can Domènech s/n, Universitat Autònoma de Barcelona; 08193 Cerdanyola del Vallès, Catalonia, Spain; Institute of Neurosciences, Universitat Autònoma de Barcelona, 08193 Cerdanyola del Vallès, Catalonia, Spain; Laboratory of Molecular Biophysics, Department of Physics, University Jaume I, 12071 Castellon, Spain

**Keywords:** Cell penetrating-peptides, membrane transport, membrane disruption, molecular dynamics simulations, cargo delivery

## Abstract

Cell-penetrating peptides (CPPs) can translocate into cells without inducing cytotoxicity. The internalization process implies several steps at different time scales ranging from microseconds to minutes. We combine adaptive Steered Molecular Dynamics (aSMD) with conventional Molecular Dynamics (cMD) to observe equilibrium and non-equilibrium states to study the early mechanisms of peptide- bilayer interaction leading to CPPs internalization. We define three membrane compositions representing bilayer sections, neutral lipids (i.e. upper leaflet), neutral lipids with cholesterol (i.e hydrophobic core), and neutral/negatively charged lipids with cholesterol (i.e. lower leaflet) to study the energy barriers and disruption mechanisms of Arg9, MAP, and TP2, representing cationic, amphiphilic, and hydrophobic CPPs, respectively. Cholesterol and negatively charged lipids increase the energetic barriers for peptide bilayer crossing. TP2 interacts with the bilayer by hydrophobic insertion, while Arg9 and MAP disrupt the bilayer by forming transient or stable pores. Collectively, these findings underscore the significance of innovative computational approaches in studying membrane-disruptive peptides, more specifically, in harnessing their potential for cell penetration.

## Introduction

The lipid fraction of biological membranes is mostly composed of phospholipids, which accounts for selective permeation, such as the cell membrane, a highly selective and dynamic barrier that encloses the contents of all living cells, responsible for cellular structural integrity, and intra- and extracellular homeostasis. Cell-penetrating peptides (CPPs) are small peptides that can be found in nature and are capable of efficiently crossing the cell membrane. CPPs optimal and efficient design to transport cargo molecules into the cell is of paramount importance ^1,2^, showing promising outcomes in fields such as, drug delivery ^3^, diagnosis of diseases ^4^, and therapeutics ^5^. According to their penetration mechanism, CPPs can translocate by passive diffusion, pore formation, endosomal escape, and endocytosis ^6^. Based on their physicochemical properties, CPPs have been classified ^7^ into cationic, such as nona-arginine (Arg9) ^8^; hydrophobic, such as Kaposi fibroblast growth factor (K-FGF) ^9^ or Translocating peptide 2 (TP2) ^10^; and amphipathic, such as Transportan 10 (TP10) ^11^ or model amphipathic peptides (MAP), a group of peptides derived from the α-helical amphipathic model peptide, designed in 1991, and here referred to as MAP ^12,13^.

From the computational perspective, translocation of any CPP is a relatively slow process and computationally too demanding to be observed in a conventional molecular dynamics (cMD) simulation ^14^. In this study, we examine the membrane disruption potential as an early step of the internalization process. We use adaptive Steered MD (aSMD) by applying an external potential followed by conventional MD (cMD) to assess whether an equilibrium has been reached (i.e. the CPPs has overcome the bilayer energy barrier to cross) or not, in order to analyse the bilayer- peptide interactions of representative CPPs such as Arg9, TP2 and MAP.

## Models and Methods

### Systems preparation

Peptides were initially modelled with ColabFold notebook ^16^, using AlphaFold ^17^ model for monomer prediction, and were relaxed in an explicit solvent system at 310.15 K. The AMBER ff14SB ^18^ force field and periodic boundary conditions were applied, and the SHAKE algorithm ^19^ was used to restrain the hydrogen atoms, allowing for a 2 fs timestep. Besides, the Monte Carlo method was used to add 150 mM KCl ions and water TIP3P molecules to solvate the system. A short minimization (5,000 cycles) and NVT equilibration (125 ps) were run with a restraint force of 1 kcal·mol^-1^·Å^-2^ on the peptide, before the unrestrained cMD simulation of 100 ns.

A peptide-bilayer system was built in CHARMM-GUI ^20–26^ for each relaxed peptide and membrane composition combination, amounting for a total of 12 systems (3 control membranes, without peptide, 1 for each bilayer, plus 9 peptide systems: 3 membrane compositions for 3 peptides). Here, peptides were placed at approximately 10 Å from the centre of mass (COM) of the upper leaflet bilayer membrane. The N-terminus or C-terminus of the peptides were not modified at any extent.

Three membrane compositions were defined: one constituted of 1,2-Dipalmitoyl phosphatidylcholine (DPPC), a neutral bilayer model commonly used in biophysical studies; a more complex one with DPPC, Dioleoyl phosphatidylcholine (DOPC) and cholesterol (CHOL), namely DPPC:DOPC:CHOL; and, thirdly, a charged membrane model containing negatively charged lipids, that is, DPPC:DOPC:DPPS:DOPS:CHOL –where DPPS stands for Dipalmitoyl phosphatidylserine and DOPS for Dioleoyl phosphatidylserine–. To avoid membrane deformation artifacts in this pulling experiment, we used 150 lipids per leaflet which, according to Hub *et al.* ^27–29^, prevents such artifacts since the bilayers are large enough. The exact composition of each membrane is the following: DPPC (150 DPPC lipids); DPPC:DOPC:CHOL (50:50:50 lipids, respectively); DPPC:DOPC:DPPS:DOPS:CHOL (30:30:30:30:30 lipids, respectively). The same conditions as in the peptide relaxing simulations were used. For the membrane lipids, the Amber Lipid21 ^30^ force field was selected.

Thereafter, the systems were minimized for 5,000 steps and equilibrated during 3.5 ns, starting in the NVT ensemble with positional restraints on the membrane atoms (restraint force of 2.5 kcal·mol^-1^·Å^-2^), and changing to the NPT ensemble after 500 ps, while lowering the positional restraints on the membrane throughout the NPT equilibration procedure (1, 0.5, 0.2, 0 kcal·mol^-1^·Å^-2^, respectively). Lastly, the membrane was relaxed for 100 ns of conventional molecular dynamics (cMD). During this step the peptide was kept restrained to avoid peptide-membrane interaction and allow for an unperturbed membrane relaxation (restraint force of 10 kcal·mol^-1^·Å^-2^).

### Adaptive steered molecular dynamics (aSMD)

Peptide translocation is a procedure computationally too expensive to observe in a conventional molecular dynamics simulation, as it commonly occurs in the scale of seconds to minutes ^14^. Consequently, we accelerated that process by using steered molecular dynamics (SMD) ^31^. SMD is a MD enhanced sampling method where an external potential is applied to accelerate the movement of a specific group of atoms -in this case, the peptide- along a defined set of coordinates. The *z* direction -the membrane normal direction- was defined as the pulling coordinate of the peptide. The reaction coordinate was defined as the distance between the COM of the carbon alpha (CA) residues of the peptide and the COM of the lipids polar head in the lower part of the bilayer, namely phosphate, nitrogen, oxygen, and the three main carbon atoms of this group.

To achieve a better approximation of the PMF and due to the large distance of the coordinate traversing the bilayer (ca. 40 Å), the membrane length (ca. 40 Å) was divided in 8 stages of 5 Å and 25 replicas were run for each step (with a constant force of 10 kcal·mol^-1^), thus using adaptive steered molecular dynamics (aSMD), as used in previous studies ^32–35^. Briefly, after each step, the Jarzynski average ^36–38^ across all replicas was calculated, and the last frame of the closest replica was used as input for the following step. Each aSMD step was run at 1 Å per ns (5 ns per replica), discussed below. An aSMD step totalled 125 ns per step and 1,000 ns per aSMD simulation. Altogether, ∼9 μs were run for the aSMD simulations of all 3 peptides.

To calibrate the system for aSMD and to determine that the membrane bilayer systems were comparable in terms of energy barrier, we performed a set of forward-backward simulations in all three bilayer systems using a single Arg residue (Arg1, Figure S1A) and, besides, we performed the simulations at two different pulling speeds (Figure S1B): 10 Å/ns and 1 Å/ns. Forward-backward PMF values for Arg1 are within the same energy interval, and at the slower velocity, the lipids had more time to adjust, leading to a lower and more accurate PMF, consistent with findings by Park, Schulten and colleagues ^39,40^. Consequently, we decided to use the slowest pulling speed for subsequent simulations, consistent with the one used by Bureau and colleagues ^41^.

### PMF Calculation

The Potential of the Mean Force (PMF) is computed by employing the Jarzynski equality ^37^ to relate the free energy difference between two states, as seen in Equation 1:

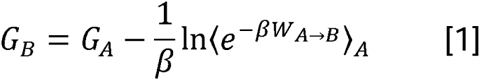

Afterwards, the replica with the closest work value to the Jarzynski average is selected as the starting point for the next simulation step.

### Conventional molecular dynamics (cMD)

Lastly, starting from the last frame of the aSMD simulation last step (where the distance between peptide and lower leaflet COMs is 0 Å), a 100 ns of unbiased cMD (also referred to as *relaxation step*) was run with the purpose of allowing the system to relax after an external potential addition. The same simulating conditions were used as in the previous cases. A total of ∼3 μs were run for the final relaxation part, accounting for 100 ns for each of the simulations (100 ns x 3 peptides x 3 membrane compositions x 3 replicas). Besides, the 3 control systems were run following the same equilibration and production protocol.

### Data Analysis

Trajectory visual analysis was performed with Visual Molecular Dynamics (VMD) ^42^, CPPTraj, and PyTraj ^43^. PyLipID ^44^ and LiPyPhilic ^45–48^ were used to analyse the simulations. An in-house Python script was implemented to compute the pore size distribution, calculating the minimum pore size in the z axis of the membrane. This script calculates the maximum distance of the water residues per each membrane z-stack and outputs the minimum distance of all the z-stacks per each simulation frame. Matplotlib ^49^ and Seaborn ^50^ were used for graphics plotting and UCSF ChimeraX ^51,52^ for molecular graphics. For the analyses, only the last 80 ns of the cMD simulation were taken into account.

## Data availability

Code to reproduce the analysis performed here is available at: https://github.com/APMlab-memb/CPPs_aSMD_cMD.git

Due to file size limitations, the simulation trajectory files will be shared upon request.

## Results and discussion

### Bilayer resistance to steered peptide crossing

The simulation protocol includes two sets of simulations: aSMD for 40 ns divided in 8 steps and 25 replicas per step to move the peptide across the bilayer defining a non-equilibrium state, followed by 3 replicas of the relaxation step, consisting in 100 ns of cMD each. This experimental design was applied to investigate the behaviour of 3 control CPPs (Arg9, MAP, and TP2) in 3 different membranes (Figure 1A). As a simplification of a complex cellular bilayer, the DPPC system is equivalent to the extracellular leaflet, richer in neutral polar headgroups, the first step for CPP internalization. DPPC:DOPC:CHOL system should equal to the second step, reaching the hydrocarbon core of the bilayer. The third step, DPPC:DOPC:DPPS:DOPS:CHOL, would be the transition from the hydrophobic core to the lower leaflet, richer in negatively charged polar headgroups.

**Figure 1.**
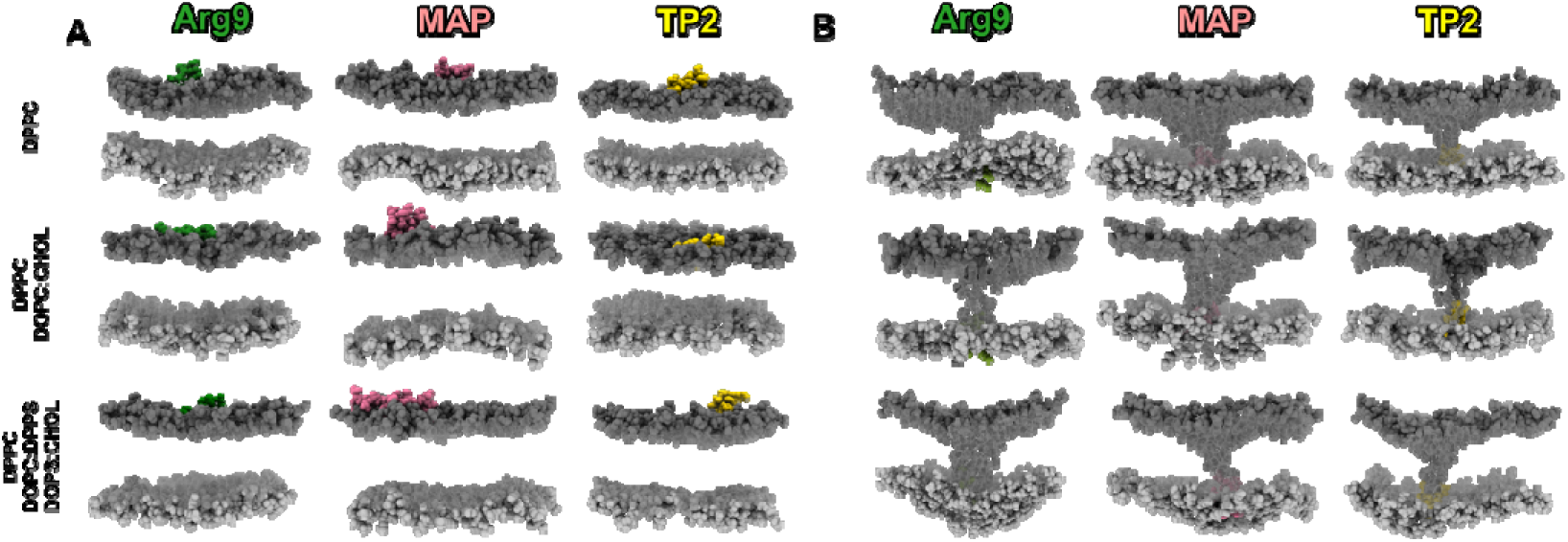
Initial and final snapshots of the aSMD process. Starting (A) and final (B) snapshots of the aSMD for the three CPPs and the three membrane compositions. See Video S1 for further detail. PMF values are indicative of the resistance opposed by the bilayer during the peptide crossing, showing that bilayer complexity is, on average, positively correlated with higher PMF values (Figure 2). In the DPPC membrane, peptides exhibit, on average, the lowest energy requirement to traverse the bilayer, indicated by a mean PMF of 181.52 ± 20.33 kcal·mol^-1^. The introduction of cholesterol to the membrane results in an overall increase in mean PMF to 200.91 ± 13.87 kcal·mol^-1^. Cholesterol has been associated with reduced efficiency in CPP translocation, –a phenomenon previously discussed by Pae *et al.* ^54^. Addition of unsaturated fatty acids (DOPC) should enhance the internalization of CPPs and lower the PMF ^55^, but this effect seems to be insufficient to compensate the influence of cholesterol. Finally, in the DPPC:DOPC:DPPS:DOPS:CHOL membrane, we observe the highest resistance to bilayer crossing, with a mean PMF of 225.65 ± 17.40 kcal·mol^-1^. Taking into consideration all three CPPs and the three bilayer systems (Figure 2), TP2 and Arg9 partition more efficiently in the upper leafltet (DPPC) compared to MAP. The transition energy from the water-bilayer interface to the hydrophobic core (DPPC to DPPC:DOPC:CHOL) is lower for MAP and TP2, and slightly higher for Arg9. Finally, from the hydrophobic core to the lower leaflet (DPPC:DOPC:DPPS:DOPS:CHOL) all peptides require higher energy for this transition, especially MAP.

**Figure 2.**
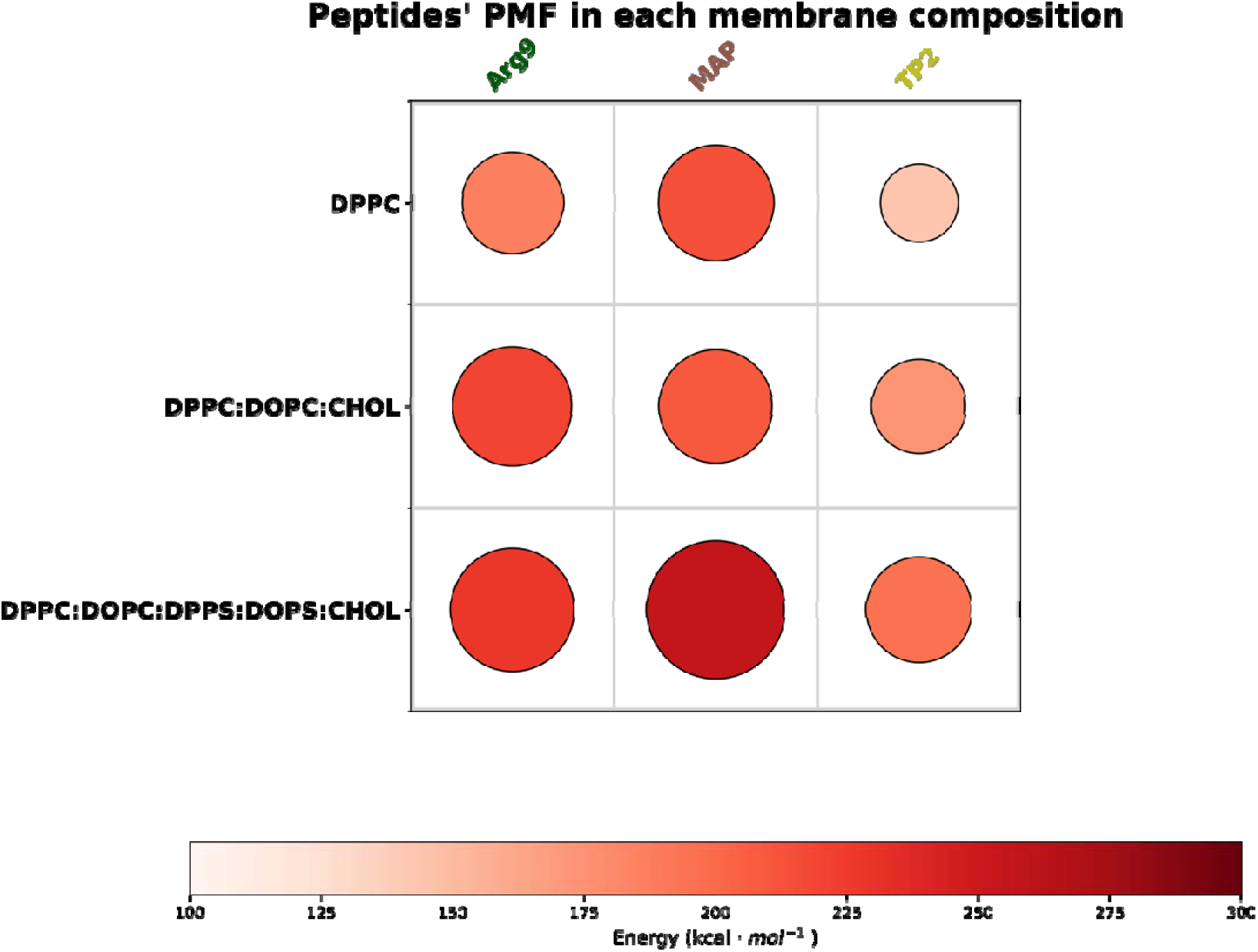
Potential of Mean Force (PMF) of peptides with respect to the membrane composition. Size and colour indicate energy. The values indicated correspond to the last value (highest energy) of the PMF analysis. PMF profiles and the PMF of all the replicas are shown in Figure S2.

After the aSMD simulation, the molecular distribution is similar for all cases (Figure 1B): the peptide has been steered into the lower part of the bilayer and is in contact with the polar heads of the lipids in the lower part of the bilayer. Some polar heads of the upper leaflet have been dragged along with the peptide during the steering process, in agreement with the previously described “Defect Assisted by Charge” (DAC) phenomenon ^53^, and the polar heads of the upper bilayer contact those of the lower bilayer.

aSMD has demonstrated PMF value accuracy calculation for peptides ^33,56–58^ and the relative trends shown for the peptides studied here are qualitatively coherent and considered as a measure to compare each peptide in the three bilayer compositions. This is of paramount importance in CPPs, where sequences differ significantly in amino acid composition, secondary structure propensities, length and physicochemical properties. Thus, quantitative assessment of PMF values should be interpreted with caution. For absolute quantitative output, computationally demanding methods with higher sampling such as, multi branched aSMD (MB- ASMD), full-relaxation aSMD (FR-ASMD) ^33^ or adaptively biasing MD (ABMD) ^32^, should be considered to obtain fully converging PMF profiles ^59^, but still, the different nature among peptides should pose a limitation.

### Peptide release after aSMD

At the end of the aSMD simulations, the peptide has been successfully transferred to the lower region of the lipid bilayer. It is important to determine whether this steered process has overcome the bilayer energy barrier reaching an equilibrium state (the energy of the process has been released) or not (the energy of the process is stored in the last step of the aSMD simulation). Thus, we performed three replicas (all with the same outcome) of cMD simulations relaxing the system to compare the peptides behaviour in each bilayer system (Video S1). At this stage we observed four possible behaviours for the peptides: (1) “Lower leaflet equilibrium state”: after the aSMD simulation, the peptide has reached an energy minimum and stays at the lower part of the bilayer; (2) “Pore formation”: the energy stored in the process results in the peptide bouncing back towards the upper leaflet remaining in the hydrophobic core and leading to formation of pores of different radius in the membrane -we define a pore as a large defect in the membrane that allows for a continuous water flow between the upper and lower leaflets; (3) “Insertion”: the energy stored in the process results in the peptide bouncing back towards the upper leaflet remaining in the hydrophobic core of the bilayer without leading to pore formation; (4) “Return”: the energy stored in the process results in the peptide bouncing back to the upper part of the bilayer. For sake of clarity, a summary of these behaviours, observed across all peptides and membrane compositions, is presented in Figure 3 and Table 2.

**Figure 3.**
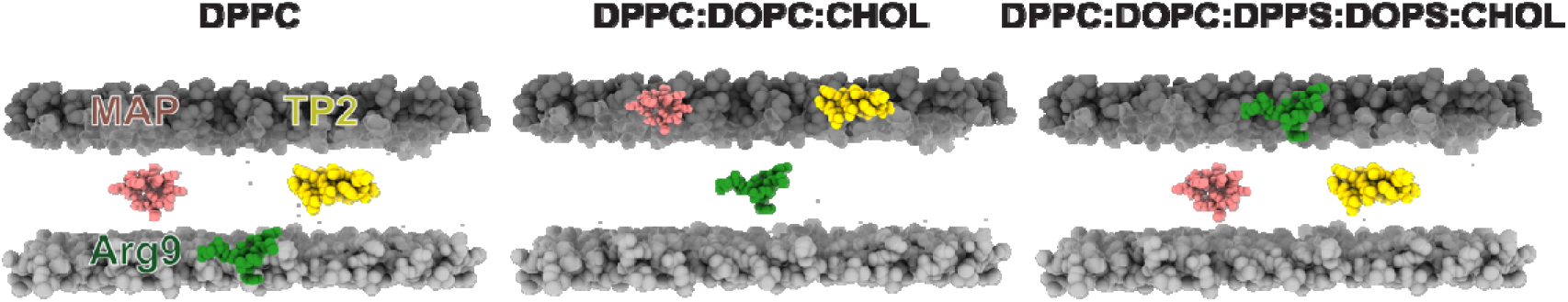
Illustrative representation of the peptide location in the 3 membrane compositions after the 100 ns of conventional MD (relaxation). Peptides are coloured as: Arg9 in dark green, MAP in rose, TP2 in gold. The polar heads of phospholipids in both the upper and lower bilayers are illustrated in darker and lighter shades of grey, respectively, while the lipid tails are portrayed in transparent white. Peptide colours are maintained in the following figures. Waters are omitted for clarity. See Video S1 for further detail.

**Table 1.**
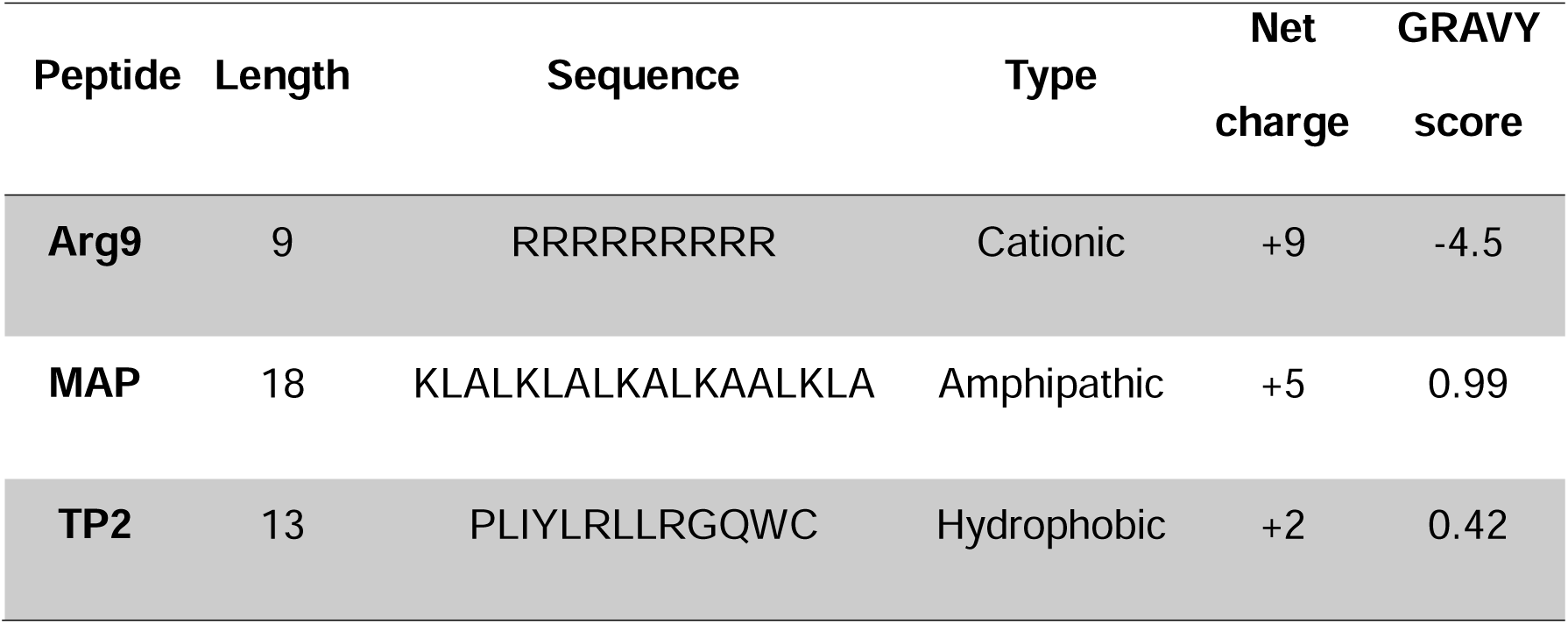
Characteristics of the peptides used in this study. GRAVY score is calculated from. ^15^.

**Table 2.**
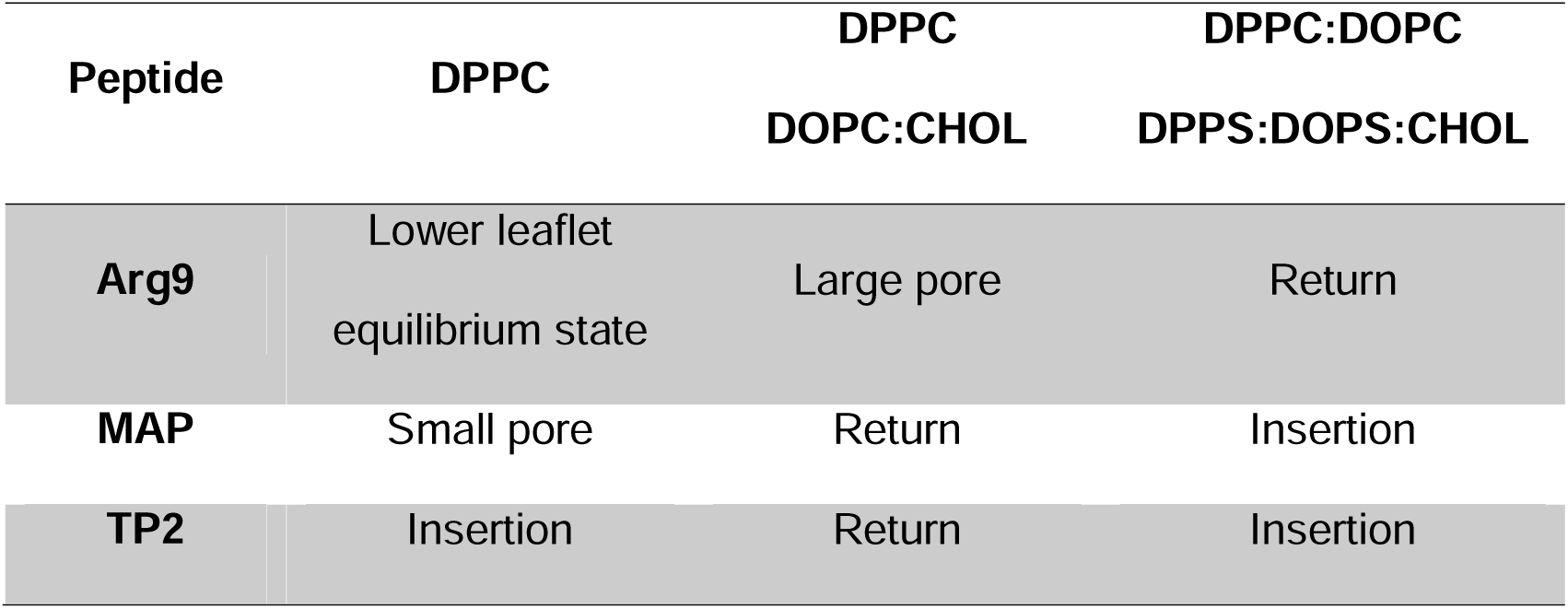
Simulation results for all CPPs in the 3 membrane compositions. All replicas show the same behaviour, and the ratios are thus omitted for clarity. Small Pore or large pore (see Table 3 for details).

| Peptide | DPPC | DPPC | DPPC:DOPC |
| --- | --- | --- | --- |
|  |  | DOPC:CHOL | DPPS:DOPS:CHOL |
| Arg9 | Lower leaflet<br>equilibrium state | Large pore | Return |
| MAP | Small pore | Return | Insertion |
| TP2 | Insertion | Return | Insertion |

Arg9 in DPPC reaches the lower leaflet after the aSMD simulation with a similar PMF value (∼200 kcal·mol-1, Figure S2) to previously published data ^32^. Besides, the energy required to move Arg9 into the middle of the membrane (from 40 to 20 Å), is similar to the energy obtained in a previous study using neutral lipids ^60^. Arg9 overcomes the imposed DPPC bilayer energy barrier since it stays in the lower leaflet for the whole cMD simulation (equilibrium state), although the formation of a small transient pore is observed (Table 3). Arg9 in the DPPC:DOPC:CHOL bilayer shows a similar behaviour in aSMD (PMF values of ∼220 kcal·mol^-1^, Figure S2) compared to previously published data ^32^. The cMD simulation for Arg9 in DPPC:DOPC:CHOL shows a relaxation from a non-equilibrium state to a more stable state where Arg9 remains trapped in the bilayer hydrophobic core while forming a large-sized pore (Table 3 and Figure S3 for pore details). In the DPPC:DOPC:DPPS:DOPS:CHOL membrane, the energy stored at the end of the Arg9 aSMD simulation is sufficient to take the peptide back to the upper leaflet.

**Table 3.** Mean radius size (Å) of the last 80 ns of the relaxation.

| Peptide | DPPC | DPPC:DOPC:CHOL | DPPC:DOPC:DPPS:DOPS:CHOL |
| --- | --- | --- | --- |
| **Arg9** | 0.19 ± 0.03 | 6.30 ± 0.04 | 0 |
| **MAP** | 0.71 ± 0.03 | 0 | 0 |
| **TP2** | 0 | 0 | 0 |

The cMD simulation for MAP in DPPC shows the peptide bouncing back but remaining in the hydrophobic core of the bilayer forming a small pore. The cMD simulation for MAP in DPPC:DOPC:CHOL shows a relaxation of the peptide and an upper part reallocation. In the DPPC:DOPC:DPPS:DOPS:CHOL bilayer, the cMD simulation for MAP shows how the peptide returns to the upper bilayer but becomes inserted in the hydrophobic core. In average, MAP has the highest PMF values, indicating that an internalization process is not as favourable as in the other cases. This can be related to experiments where they observed that the internalization of MAP requires, in a large amount, an energy-dependent pathway or vesicle transport event ^12,61–63^.

Similarly to MAP, TP2 has not reached an equilibrium in the lower part of the bilayer in any condition. In DPPC:DOPC:CHOL, TP2 releases all the stored energy and returns to the upper bilayer, indicating that cholesterol induced rigidity poses a high energy barrier for TP2 to remain in the bilayer. On the other hand, in DPPC and DPPC:DOPC:DPPS:DOPS:CHOL bilayers, we observe the insertion of TP2 in the hydrophobic core of the bilayer, but without leading to the formation of a pore. This behaviour can be related to the fact that TP2 in monomeric form enters the cell via spontaneous membrane translocation, rather than the pore formation mechanism64,65

Effects of the peptides on bilayer behaviour have been performed, namely lipid order parameter and membrane thickness (Figure S3). All membranes show a similar thickness (around 40 Å), with the DPPC membranes showing a smaller value, since the addition of cholesterol causes an increase in membrane packing and, consequently, increased membrane thickness ^66^. Lipid order parameter analyses show that membranes are well organized, and no significant differences are observed between membranes regardless of the type of peptide-bilayer interaction. Thus, we have focused on the occupancy of peptide residues by the polar heads of the phospholipids in upper and lower leaflets (Figure 4).

**Figure 4.**
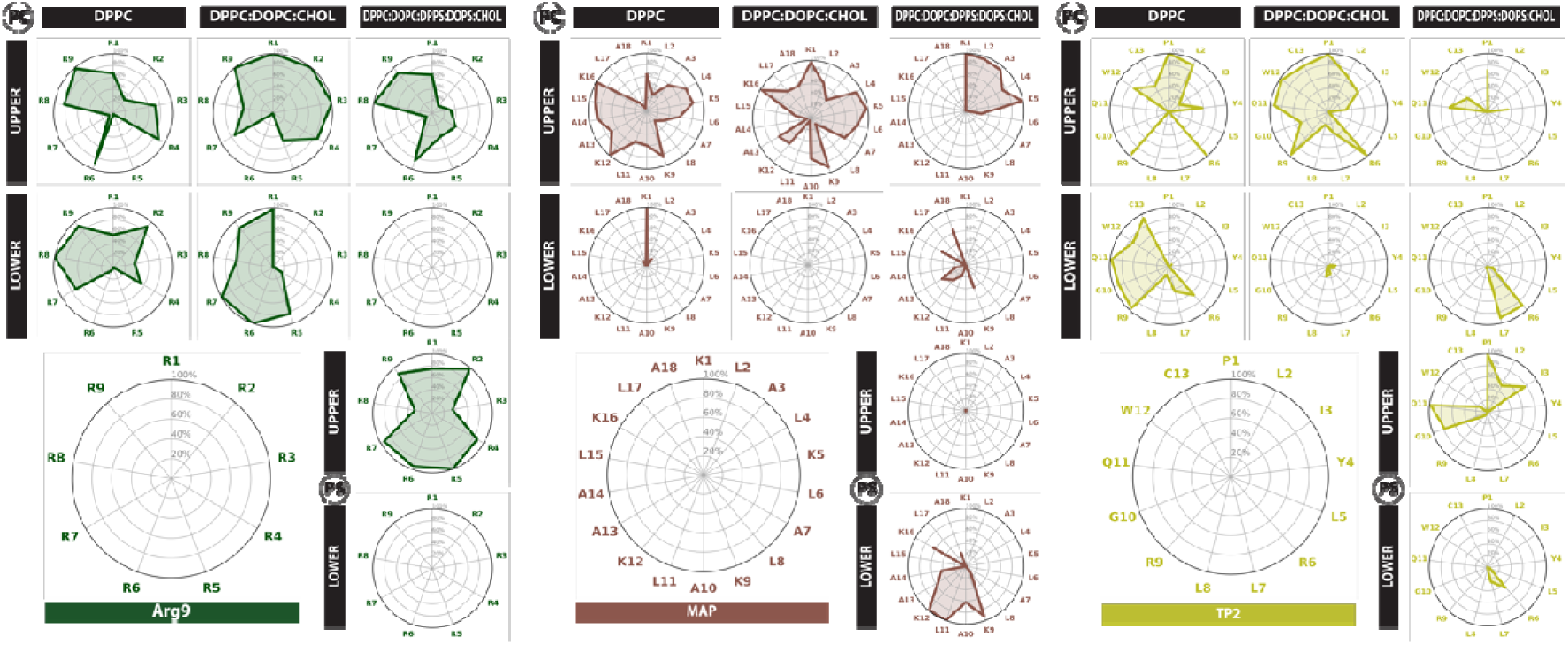
CPPs’ residue percentage occupancy by the polar head of the phospholipids in the upper and lower leaflets. Polar heads corresponding to PC (phosphatidylcholine) occupancy is shown for DPPC, DPPC:DOPC:CHOL and DPPC:DOPC:DPPS:DOPSCHOL membranes. For the third membrane, PS (phosphatidylserine) occupancy is also shown. The occupancy analysis refers to the first replica.

### Peptide-bilayer interactions

Sequence composition, charge and hydrophobicity (Table 1) are key determinants driving peptide-bilayer interactions. Arg9 has a high positive charge and hydrophilicity. Arg9 shows 3 different modes: lower bilayer steady-state, upper part relocation or pore formation, but, in each of these, it stays in contact with the polar heads of the bilayer (in the two former cases) or with the waters (in the latter case), demonstrating that it prefers to interact with hydrophilic interfaces. In the other two canonical CPPs, we see how they get inserted into the bilayer and stay in contact with the hydrophobic part of the membrane, which is related to the more hydrophobic nature indicated by the GRAVY score. In parallel, both MAP and TP2 have key positively charged residues (Table 1), which allows them to interact with the polar heads in the membrane.

In Figure 4 we present the occupancy analysis regarding the lipids’ polar heads for every peptide in all membranes, such as, PC for phosphatidylcholine (in DPPC/DOPC) and PS for phosphatidylserine (in DPPS/DOPS). Occupancy is defined as the percentage of simulation time that a residue is in contact with a lipid. In Figure S4, we show the occupancy by the lipid tails and cholesterol. Regarding peptide-polar head interactions (Figure 4), we observe a higher interaction ratio for Arg9 (several residues have 100% occupancy), which can be explained due to the polycationic nature of this CPP, strongly attracted to the negatively charged polar heads of the lipids. K/R neighbouring residues also show high occupancy in all three CPPs. MAP, which has alternating positive (K) and hydrophobic (L, A) residues, preferably interacts with the polar heads through positive residues, that is, K1, K5, K9, K12, and K16. TP2 contains only two charged residues, R6 and R9, which are prone to interact with the polar heads of the lipids and show high occupancy across the three bilayers. However, the N- and C-terminal parts are also interacting with the polar heads in three and two bilayers, respectively. In DPPC, the peptide is inserted in the membrane and stretched, thus interacting with a leaflet in each end. In the DPPC:DOPC:CHOL bilayer, R9 favours the lipid interaction of TP2 C-terminal residues. Besides, the N-terminal residues (specially P1) show high occupancy, which can be explained by the positive charge in the N-terminal residue. On the other hand, regarding the occupancies by lipid tails, Arg9 shows, in average, low occupancy, again, explained by its polycationic nature, whereas MAP and TP2 show high occupancy by the lipid tails, mainly interacting with the hydrophobic residues (L, A in MAP, and L, I, Y, W in TP2).

In parallel, when comparing the occupancies across all three bilayers, there are noteworthy differences between: (1) the case where the peptide that has reached an equilibrium in the lower part of the bilayer, which has a higher occupancy in the lower leaflet (Arg9 in DPPC), (2) the peptides that form a pore and interact with the polar heads in upper and lower leaflets (MAP in DPPC, and Arg9 in DPPC:DOPC:CHOL), (3) the peptides that get inserted into the bilayer and also interact with both leaflets (TP2 in DPPC, MAP and TP2 in DPPC:DOPC:DPPS:DOPS:CHOL), and (4) the peptides that have been reallocated to the upper leaflet and are only interacting with the polar heads in the upper leaflet (MAP and TP2 in DPPC:DOPC:CHOL, Arg9 in DPPC:DOPC:DPPS:DOPS:CHOL). MAP in the first membrane composition generates a pore in the bilayer, and only interacts with the lower leaflet with the first residue, indicating that it shows an extended conformation, perpendicular to the membrane ^67,68^, stabilized by the hydrophobic interactions with the lipid tails and the hydrophilic interactions with water, with a similar distribution to the three cases of insertion (see Video S1). Interestingly, Arg9 in DPPC:DOPC:DPPS:DOPS:CHOL interacts rather with the polar heads in PS lipids than PC lipids, likely by the strong attraction between the side chains and the negatively charged lipids, as seen in previous studies ^69^.

The cationic Arg9 seem to require pore formation to cross the bilayer ^70,71^ as we observe for Arg9 by either forming transient (DPPC bilayer) or more stable (in the DPPC:DOPC:CHOL bilayer), likely a mechanism to overcome the bilayer energy barriers. MAP shows the highest resistance to bilayer crossing, which can be lowered by the formation of a pore ^72^, related to what we see in DPPC. TP2 may involve direct translocation (through a quick and transient pore or without pore formation as we observe here) of a monomeric peptide, leading to a minimum leakage.

Altogether, we see how interactions with the upper leaflet strongly influence the ability of the peptides to interact with the lower leaflet and, consequently, their ability to form pores or reach the lower leaflet. The presence of cholesterol adds more stability to the membrane and increased thickness, which entails higher deformation resistance, as seen by higher PMF values, and less events of bilayer crossing. In the most complex membrane, there are no bilayer crossing or pore-forming events, which agrees with the fact that negative lipids are not directly correlated with the internalization efficiency of CPPs (Elber, 2023).

From this study, we could argue that each peptide may find different energetic barriers at different bilayer levels. For MAP, the upper leaflet partitioning is limiting, a possible solution could be transient water pore formation to decrease this energy, as discussed before. For Arg9 the peptide-lipid and peptide-water interactions, which may lead to larger disturbance of the bilayer and formation of large transient pores would be key to overcome the energy barrier at the hydrophobic core layer. The hydrophobic TP2 seems to find the easier path across the bilayer, with lower energy barriers, without requiring the formation of transient pores. Other aspect that requires further investigation beyond the scope of the present study, that should not be ruled out regarding the internalization mechanism for CPPs, are secondary structure conversions, peptide organization and/or peptide self-assembly ^53,60,69,73,74^.

Despite the significant strides made in the understanding of CPP internalization, translocation and pore formation, certain aspects remain elusive, which underscores the need for future investigation, as well as the need for out-of-the-box ideas to study such processes.

