## Supplemental Figures for "Computational Insights into Membrane Disruption by Cell-Penetrating Peptides"

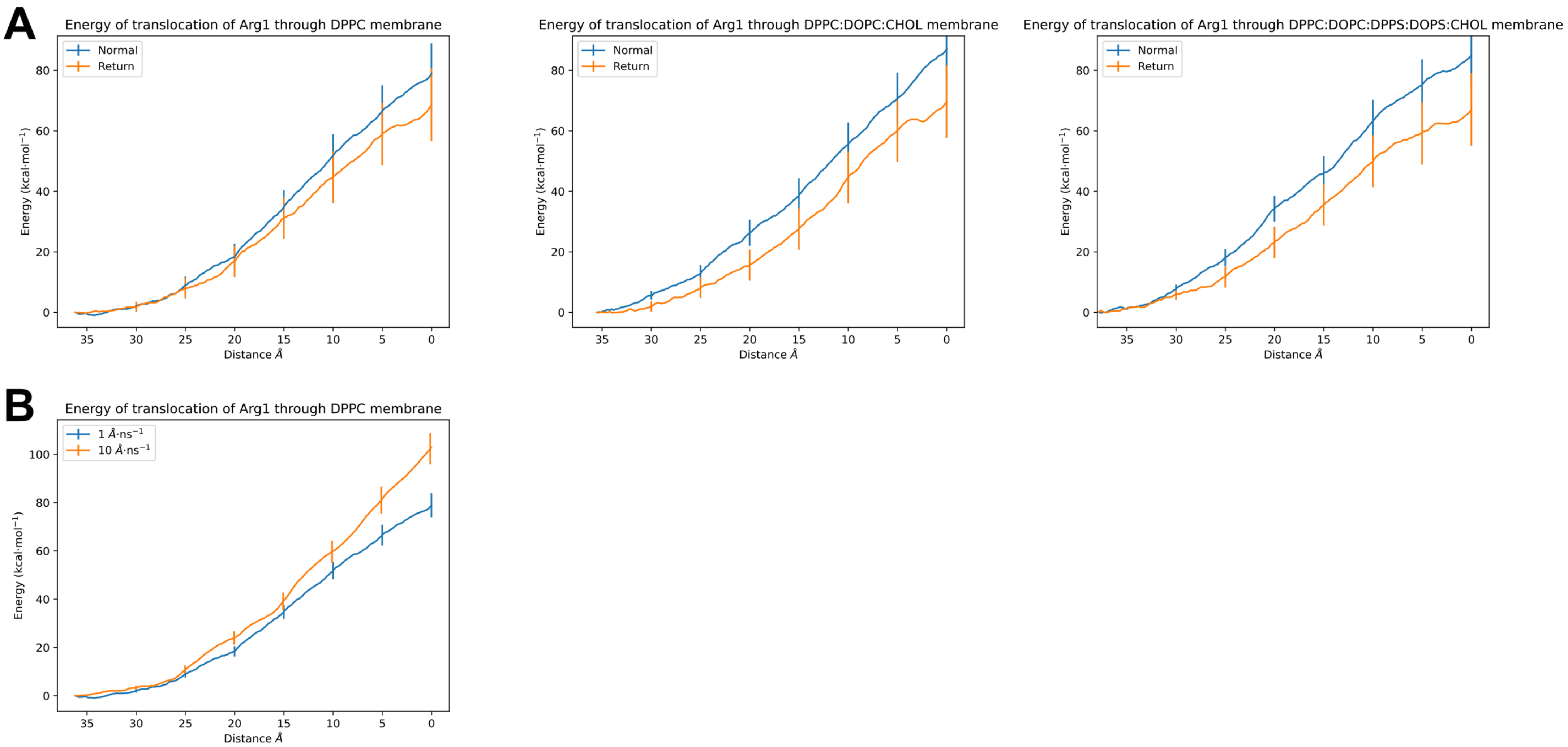


**Figure S1. PMF calculation in (A) normal and return aSMD simulations and (B) different pulling speed simulations.** (A) The effect of normal and return aSMD has been computed for the three membrane compositions. (B) The effect of pulling speed has been done in DPPC membrane.

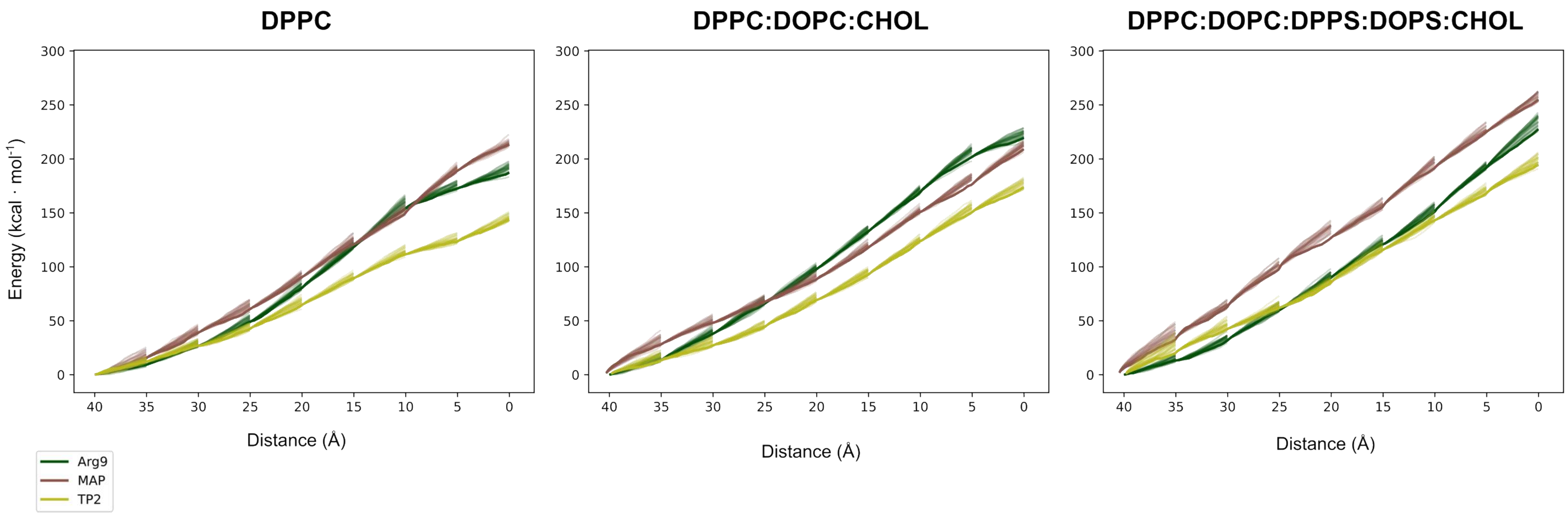


**Figure S2. PMF profiles of CPPs.** The PMF correspond to DPPC, DPPC:DOPC:CHOL, and DPPC:DOPC:DPPS:DOPS:CHOL membranes, respectively. The PMF of all the replicas are also shown, with a transparency of 20%.

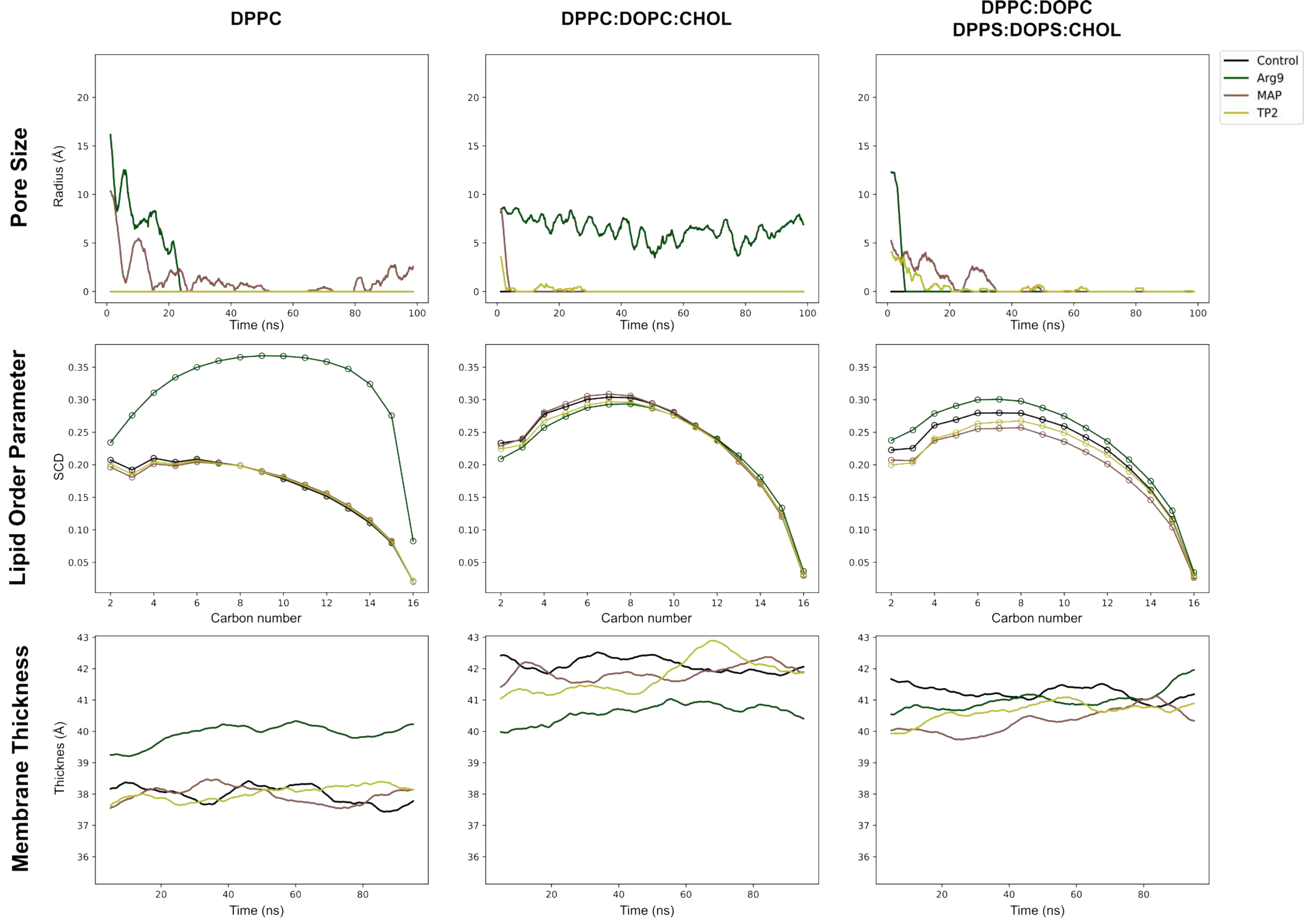


**Figure S3. Pore size, lipid order parameter and membrane thickness analyses.** The analyses have been performed in the cMD part of the DPPC, DPPC:DOPC:CHOL and DPPC:DOPC:DPPS:DOPS:CHOL membrane simulations. Lipid Order Parameter has been computed for the lipid tails and results are shown for carbon number 2 to 16.

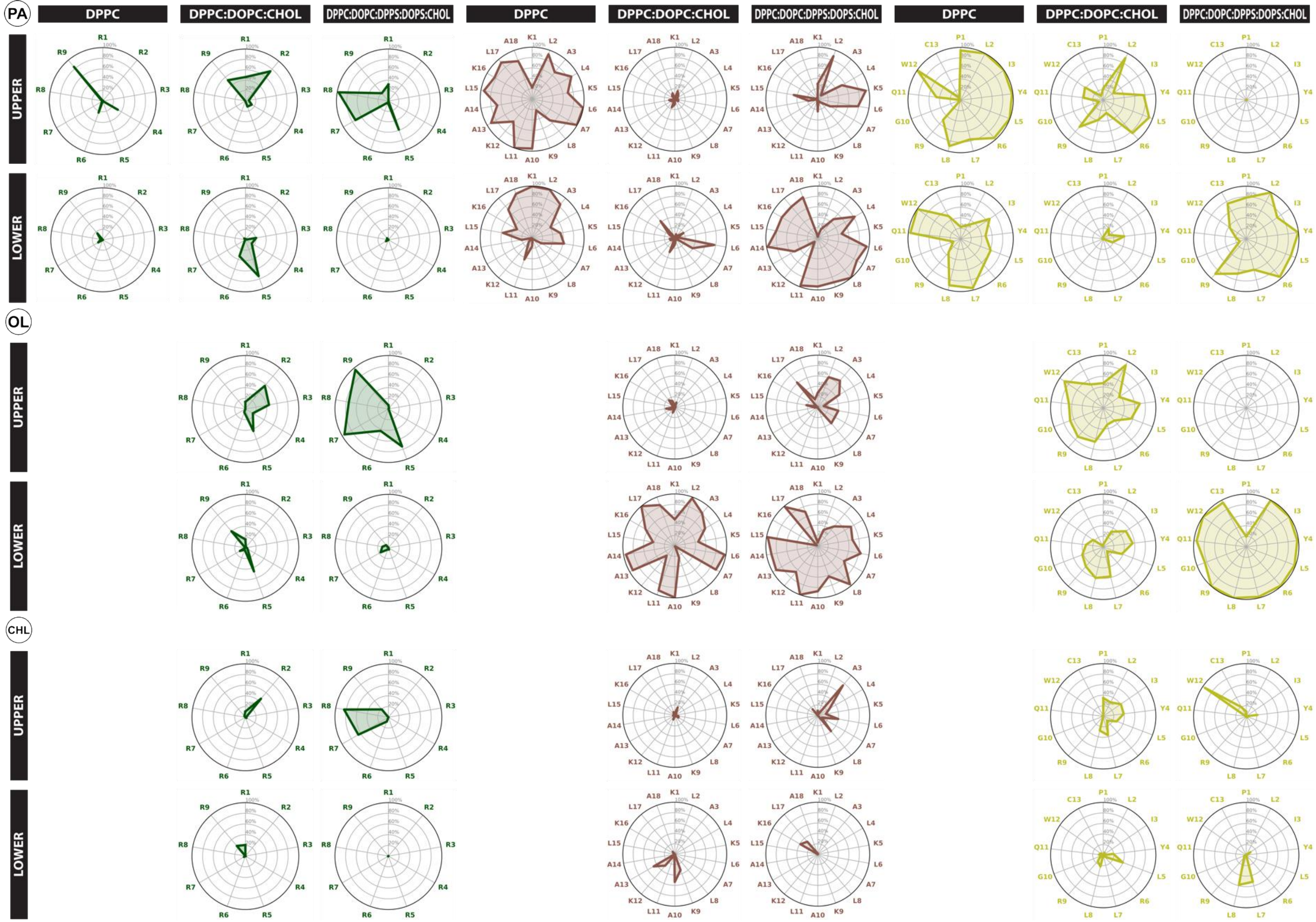


**Figure S4. Occupancy of the lipid tails and the cholesterols.** PA refers to the lipid tail present in DPPC/DPPS lipids, namely palmitic acid. OL refers to the lipid tail in DOPC/DOPS, namely oleic acid. CHL refers to cholesterol. These are the lipid names provided by the AMBERFF14SB forcefield.
